# Chronic alcohol drinking delays recovery from capsaicin- and nerve injury-induced hypersensitivity in mice

**DOI:** 10.1101/2025.09.05.674513

**Authors:** Rachel Schorn, Maureen Riedl, Laura S. Stone, Anna M. Lee, Lucy Vulchanova

## Abstract

Chronic pain and alcohol use disorder (AUD) are major health concerns that significantly impair quality of life. Persistent pain is common in individuals with alcohol dependence, and alcohol is commonly used by chronic pain patients for self-medication. The neural mechanisms linking these conditions remain unclear. We hypothesized that chronic alcohol exposure induces hypersensitivity in multiple modalities and increases the duration of recovery in acute and persistent pain models. Using the two-bottle free-choice alcohol consumption paradigm in adult mice, alcohol-induced hypersensitivity (AIH), indicated by reduced mechanical withdrawal thresholds, developed after 4-6 weeks of alcohol consumption compared to age- and sex-matched water-consuming controls. Alcohol-consuming female mice, but not male mice, developed cold hypersensitivity after AIH emerged. To assess the impact of chronic alcohol consumption on acute and persistent pain, we used intraplantar capsaicin and sciatic nerve crush models, respectively. In the capsaicin model, water-treated, but not alcohol-treated, mice recovered from hypersensitivity by 24 hours post-injection. In the sciatic nerve crush model, alcohol-consuming mice exhibited slower recovery of mechanical withdrawal thresholds compared to water-consuming controls. While mechanical hypersensitivity in water-consuming mice returned to pre-surgical thresholds by 3-4 weeks post-surgery, recovery in alcohol-consuming mice was both delayed and partial. Surgical intervention did not impact alcohol intake. Overall, our results suggest that chronic voluntary alcohol consumption facilitates the transition to chronic pain by prolonging hypersensitivity and delaying recovery from injuries.

**Highlights:**

- Magnitude of alcohol-induced mechanical hypersensitivity is similar across sex
- Females, but not males, develop alcohol-induced cold sensitivity
- Chronic alcohol prolongs acute inflammatory pain
- Chronic alcohol prolongs nerve-injury induced neuropathic pain

## 1. Introduction

Alcohol use disorder (AUD) is characterized by mild to severe symptoms of problematic alcohol use despite adverse social, occupational, or health consequences. The COVID-19 pandemic exacerbated problematic alcohol use in both adults and adolescents [1–3]. In the United States, 28 million people ages 12 or older had AUD in 2024 [4].

Chronic pain and AUD are often comorbid [5] and this comorbidity is bidirectional, creating a cycle of escalating alcohol use and worsening pain. Individuals with chronic pain are significantly more likely to self-medicate with alcohol to manage pain symptoms compared to those without chronic pain [6–11], particularly when drinking to intoxication where blood alcohol concentration (BAC) exceeds 0.08% or 80 mg/dL [12, 13]. Additionally, approximately 30-40% of individuals with alcohol use disorder (AUD) report experiencing chronic pain lasting 3 months or longer [14, 15]. AUD increases the likelihood of developing chronic pain [13, 14], suggesting that chronic alcohol use contributes to pain persistence. Moreover, pain is a risk factor for relapse for AUD patients [16], and evidence suggests that effective pain management reduces the risk of relapse [17]. The present study emphasizes AUD as a precursor to chronic pain to investigate how alcohol affects pain recovery and whether it contributes to long-term hypersensitivity. Studies investigating how chronic alcohol alters pain processing are critical for optimizing effective pain management during alcohol cessation treatment to break the cycle of comorbid chronic pain and AUD.

The interactions of chronic pain and voluntary chronic alcohol remain poorly understood at a mechanistic level, particularly regarding long-term effects on the central nervous system. Studying this comorbidity in humans presents significant ethical and practical challenges, including controlling for variables such as alcohol intake, pain severity, and adherence to treatment over long periods, which leads to difficulties in establishing causation. To bridge this gap, mouse models offer a controlled approach to address these challenges. There is extensive literature on rodent models of AUD, and increased sensitivity to noxious thermal and mechanical stimuli has been consistently observed in a variety of paradigms. Withdrawal-induced hypersensitivity has been observed during protracted withdrawal following cessation of chronic alcohol exposure and during repeated acute withdrawal in intermittent exposure paradigms involving adult and adolescent mice [18–25]. Many models use non-voluntary alcohol exposure via oral gavage or vapor inhalation. While this allows for more precise alcohol dosing, involuntary access paradigms fail to mimic naturalistic human drinking behavior. We therefore utilized a voluntary intermittent access paradigm termed chronic intermittent alcohol two-bottle choice (CIA-2BC). The CIA-2BC model allows for observation of acute and chronic effects of alcohol consumption [21–25]. This model is advantageous for examining how voluntary alcohol consumption interacts with persistent pain, as it replicates the cycle of repeated drinking and withdrawal seen in individuals with AUD.

Acute alcohol administration is analgesic in inflammatory and neuropathic pain models [26, 27], while chronic alcohol administration is thought to be pronociceptive [25, 28, 29]. Notably, the effects of chronic alcohol consumption on the transition to persistent pain are largely unexplored. To model the effect of chronic alcohol consumption on acute inflammatory pain, we measured mechanical hypersensitivity following intraplantar capsaicin injection. Capsaicin induces acute localized pain and inflammation that resolves in 24-48 hours. To model the effect on persistent pain, we used the sciatic nerve crush injury. This model reliably induces transient behavioral signs of neuropathic pain, including mechanical hypersensitivity, that typically last 3-4 weeks. Neuropathic pain is experienced by humans with comorbid chronic pain and AUD [30, 31]. Combining the CIA-2BC paradigm with the nerve crush model allowed us to study the bidirectional influence of chronic alcohol use on persistent pain. We hypothesized that chronic voluntary alcohol consumption induces hypersensitivity and delays recovery from capsaicin- and nerve injury-induced hypersensitivity.

## 2. Methods

### 2.1 Animals

Adult age-matched male and female C57BL/6J mice aged 8-12 weeks upon the start of alcohol consumption were used in this study. C57BL/6J mice were used given their established preference for alcohol consumption compared to other inbred strains of mice [32, 33] and their frequent use in both alcohol and pain pre-clinical studies [19, 34–38]. Due to the large number of mice required for this study, animals were processed in multiple sub-cohorts, with each treatment group represented within each sub-cohort to mitigate against any batch effects. Figures 2 through 5 contain data from cohort 1, while figures 6 & 7 include data from cohort 2. Vivarium rooms were temperature (∼21 C) and humidity (∼70%) controlled on a 12-hour light/dark cycle. Animals were housed in conventional cages with access to food and water ad libitum. Procedures and experimental protocols were compliant with the Guide for the Care and Use of Laboratory Animals were approved by the University of Minnesota Institutional Animal Care and Use Committee.

### 2.2 Voluntary CIA consumption paradigm

To investigate the effects of voluntary chronic alcohol consumption, chronic intermittent alcohol 2-bottle choice (CIA-2BC) was implemented for 9 weeks (**fig. 1**). The procedure was adapted from prior work [37, 39]. Mice were individually housed to monitor alcohol intake. The drinking protocol consisted of alternating 24-hour periods of alcohol and water access, with alcohol available on Mondays, Wednesdays, and Fridays. On alcohol days, mice had access to two bottles in their home cage: one containing 20% alcohol (v/v) and the other containing water. On water-only days, mice had access to two water bottles. The first week of alcohol exposure involved gradually increasing alcohol concentrations (3% on Monday, 10% on Wednesday, 20% v/v on Friday). Alcohol concentration was 20% v/v on all subsequent weeks. Body weights were measured weekly. Alcohol consumption is reported as the amount consumed relative to body weight ((weekly average g of alcohol solution consumed over 24h × alcohol density 0.789 × alcohol concentration)/mouse body weight in kg) and as percent preference for the alcohol bottle (weekly average of daily preferences: (g of alcohol solution consumed over 24h) / (g of alcohol solution + g of water consumed over 24h) × 100). To account for leakage from bottles during home cage consumption, bottles were administered to an uninhabited control cage Mondays, Wednesdays, and Fridays, and weighed alongside the alcohol and water bottles.

**Figure 1.**
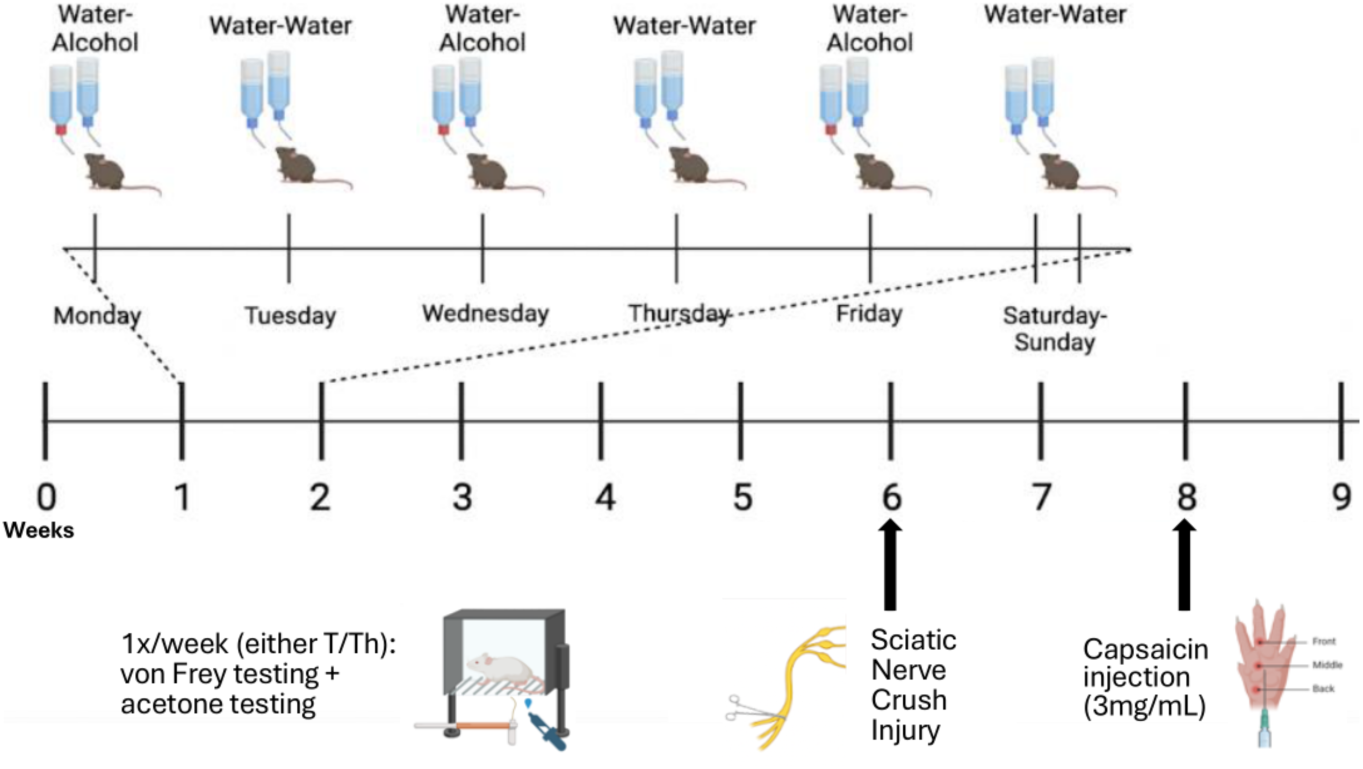
Diagram of the timeline of voluntary alcohol consumption using the intermittent access two-bottle choice model. Mechanical thresholds were measured on Tuesdays or Thursdays (water only days) using von Frey filaments and computed using the up down method. In some cases, cold sensitivity was assessed on the same days immediately following von Frey testing. Sciatic nerve crush was performed immediately after von Frey testing during week 6, and capsaicin injections were performed immediately after week 8 testing. Figure created with BioRender.com.

### 2.3 Mechanical sensitivity testing

To assess mechanical hypersensitivity, von Frey monofilaments were used according to the Up-Down Reader method [40]. This method is based on the application of monofilaments to the plantar surface of both hind paws, with increasing force until a paw withdrawal response occurs. Mice were tested weekly or every two weeks on a water-only day (either Tuesdays or Thursdays) for the duration of the experiment by the same female experimenter.

### 2.4 Cold sensitivity testing

Cold sensitivity was assessed immediately after completion of von Frey testing using previously reported methods [41] (**fig. 1**). To measure cold hypersensitivity, 25 µL of acetone was administered to the left hind paw using a fixed volume micropipette (United Scientific Fixed Volume Mini Pipettes #USCI-PMP025). Videos of the mouse’s ventral surface were recorded for one minute after acetone application using a webcam (Logitech StreamCam #960-001286). Responses specific to the treated paw, including lifting, licking, shaking, and scratching, were manually scored from each video by measuring response duration in seconds.

### 2.5 Acute inflammatory pain model (capsaicin injection)

To produce inflammation-induced hypersensitivity, a unilateral hind paw injection of capsaicin was performed after 8 weeks of drinking (**fig. 1**). After anesthetizing with 2.5% isoflurane vapor, ophthalmic ointment was applied to the eyes and the mouse’s left hind paw was disinfected with betadine. One 10 µL injection of capsaicin (3 mg/ml capsaicin + 10% DMSO in saline) or vehicle (10% DMSO in saline) was given intradermally into the left hind paw using a 50 µL Hamilton syringe and 30-gauge needle.

### 2.6 Persistent pain model (sciatic nerve crush)

To induce nerve injury-induced hypersensitivity, a sciatic nerve crush surgery was performed after week 6 von Frey testing according to methods described in [42] (**fig. 1**). After anesthetizing with 2.5% isoflurane vapor, ophthalmic ointment applied to the eyes and the mouse’s left hindquarters were prepared by shaving with surgical clippers and cleansing with betadine. The mice were positioned on a heat pad to maintain a normal body temperature. The leg was elevated on a gauze pad and secured with tape. The sciatic nerve crush procedure involved a midline incision and dissection of the hamstring muscle, and exposure of the sciatic nerve. Once the three branches of the nerve were visualized, super-fine hemostatic forceps (Fine Science Tools #13020-12) was used to crush the entire diameter of the nerve for 10 seconds. To standardize compression pressure, the nerve was positioned ∼1mm into the forceps, and the first notch on the forceps was used. The surgical area was then sutured, and each mouse underwent IACUC-approved post-operative care. CIA-2BC continued following sciatic nerve crush.

### 2.7 Statistical analysis

Data analysis was performed using GraphPad PRISM. Data were analyzed for significance with three-way or two-way repeated measures (RM) ANOVAs with multiple comparisons tests, and simple linear regression. Results are presented as mean ± SEM (represented by error bars). A p-value of <0.05 was considered statistically significant. As there are known sex differences in alcohol consumption and preference in C57BL/6J mice [32, 43, 44] and known sex differences in alcohol-and pain-related mechanisms in C57BL/6J mice [29, 34, 35, 38, 45–47], data was analyzed with sex as a variable for specific comparisons to better delineate the interactions between alcohol and pain in each sex. For certain analyses, such as the capsaicin and nerve crush experiments, sexes were separated to simplify the statistical comparisons.

## 3. Results

### 3.1 Voluntary chronic alcohol consumption produces alcohol-induced mechanical hypersensitivity (AIH)

Consistent with prior literature, females had greater alcohol consumption (g/kg) than males at all weeks of the study (**fig. 2a**, RM ANOVA F_sexXtime_ (7,182)=1.26, *p*=0.27, F_time_ (7,182)=4.38, *p*=0.0002, F_sex_ (1,26)=51.48, *p*<0.0001). Females had a weekly average of 17.6 g/kg of alcohol consumption per 24h, while males averaged 8.4 g/kg per 24h across the duration of the study, indicating that females consumed 2.1-fold more alcohol per kg body weight compared with males. Female mice also showed increased alcohol preference compared with males across the duration of the study (**fig. 2b**, RM ANOVA F_sexXtime_ (7,182)=0.96, *p*=0.46, F_time_ (7,182)=7.32, *p*<0.0001, F_sex_ (1,26)=12.62, *p*=0.002), and had an average of 22.9% greater alcohol preference for the 20% alcohol concentration compared to males for the duration of the study. To evaluate the impact of chronic alcohol consumption on mechanical sensitivity, we measured mechanical withdrawal thresholds weekly during the drinking paradigm. The mechanical thresholds were averaged between left and right hind paws, as no significant differences were observed between sides for individual mice at any time point, indicating that the effects of alcohol consumption were bilateral (RM ANOVA, no time by side interaction, no main effect of time or main effect of side, all *p*>0.05).

**Figure 2.**
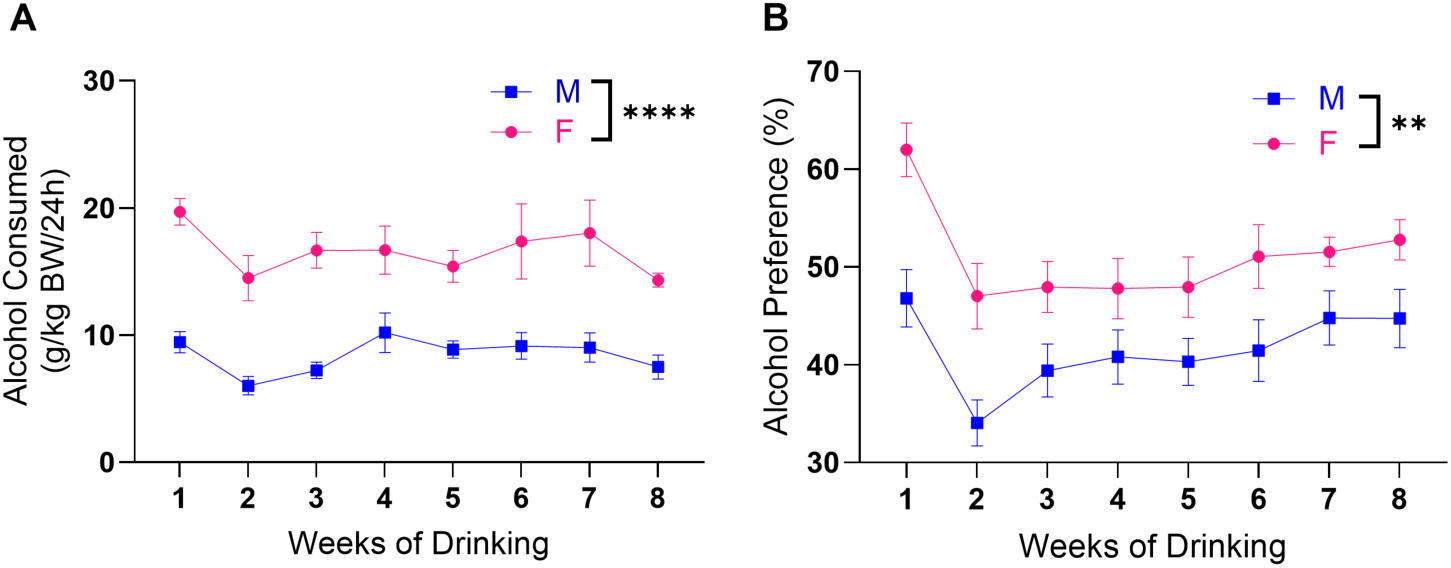
Alcohol consumption is greater in C57BL/6J females. Female mice voluntarily drink more alcohol than male mice. Average alcohol consumed per day measured relative to **(A)** body weight and **(B)** % preference for alcohol for males (*n*=15) and females (*n*=13). RM ANOVA, main effect of sex \*\**p*=0.002, \*\*\*\**p*<0.0001.

As we observed that female mice consumed on average 2.1-fold more alcohol and had 22.9% greater preference for the alcohol than the males, we were interested in whether the amount of alcohol consumed was related to changes in mechanical sensitivity. To assess mechanical AIH, the percent change in paw withdrawal threshold from baseline was calculated at each week of drinking (**fig. 3a**). The greatest magnitude of mechanical AIH was observed after 6 weeks of alcohol for males and after 5 weeks for females, where tactile thresholds decreased on average by 28.3% in males and 36.9% in females, respectively. Although the magnitude of the maximal difference in mechanical AIH was larger in females than males, it was not significantly different between sexes at any time point (**fig. 3a**, RM ANOVA, F_timeXsex_ (5,125)=0.71, *p*=0.62, F_time_ (3.14, 78.41)=31.51, *p*<0.0001, F_sex_ (1,25)=1.80, *p*=0.19). These data indicate that the degree of mechanical hypersensitivity does not correlate with the amount of alcohol consumed, and that the magnitude of the AIH was similar between males and females despite females consuming 2.1-fold more alcohol per body weight compared with the males.

**Figure 3.**
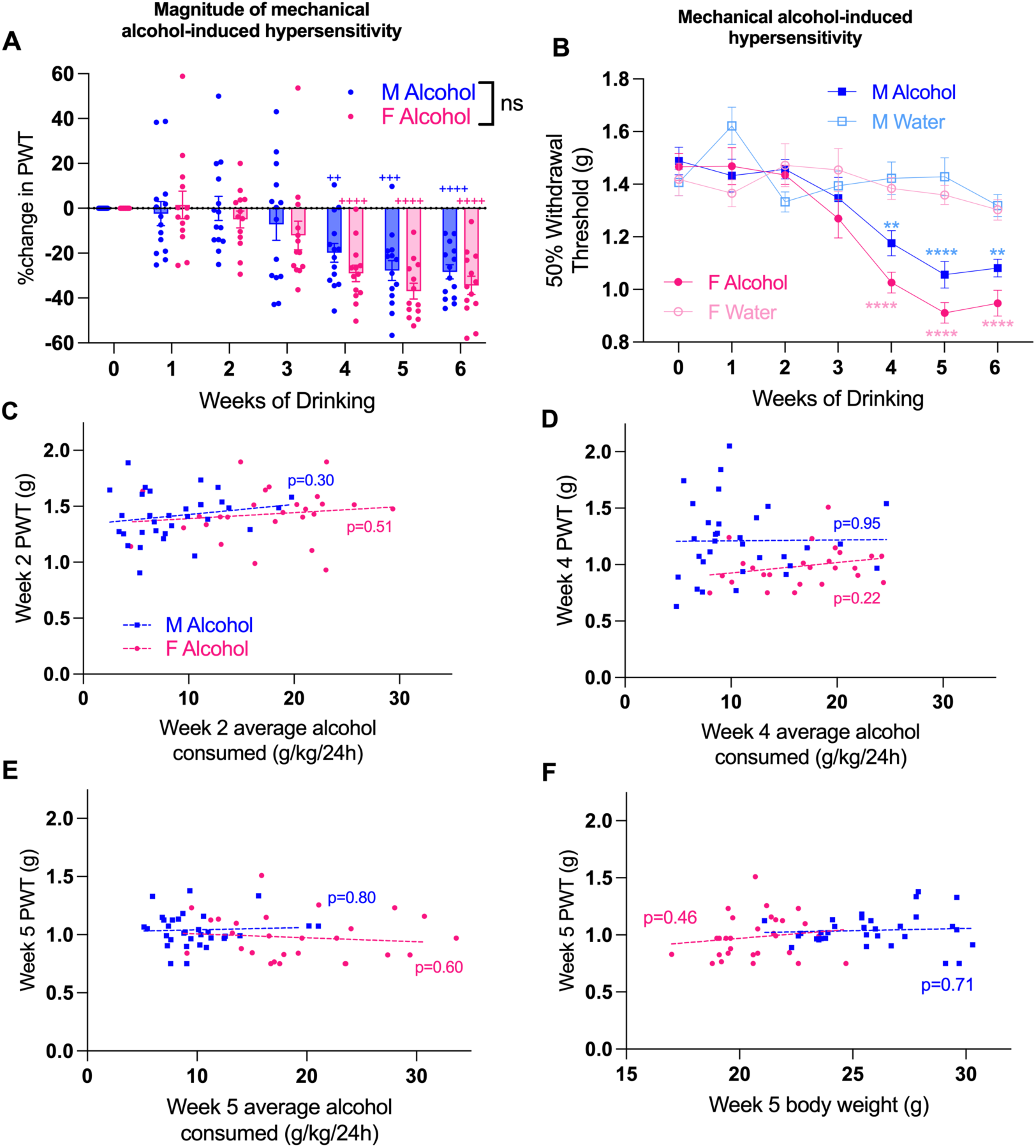
Chronic alcohol consumption produces alcohol-induced mechanical hypersensitivity (AIH). **(A)** No difference in magnitude of mechanical AIH between males and females (*n*=13-14/group/sex, Sidak’s multiple comparisons test *p*>0.05, males vs. females weeks 4-6), determined by changes in paw withdrawal thresholds (PWT). Significant reductions in PWT (relative to week 0 baseline) appeared from 4-6 weeks of alcohol consumption (Sidak’s post hoc test, ^++^*p*<0.01, ^+++^*p*<0.001, ^++++^*p*<0.0001). **(B)** Alcohol-consuming mice developed mechanical hypersensitivity 4-6 weeks after initiation of alcohol exposure (*n*=13-14/group/sex; Tukey’s multiple comparisons test \*\**p*<0.01, \*\*\*\**p*<0.0001 relative to same-sex water-drinking mice). Hind paw thresholds were averaged, since there was no significant difference between left/right thresholds within any group during weeks 1-6 (Tukey’s multiple comparisons test *p*>0.05). **(C-E)** Simple linear regression analyses of paw withdrawal threshold vs amount of alcohol consumed in male (*n*=32) and female (*n*=26) mice following 2-, 4-and 5-weeks of alcohol consumption (C, D and E, respectively) indicates paw withdrawal thresholds are not directly proportional to alcohol consumption (*p*>0.05). **(F)** No relationship was observed between body weight and paw withdrawal threshold at 5 weeks.

We then examined the 50% paw withdrawal threshold each week and also found no significant sex differences in mechanical sensitivity between alcohol drinking males and females at any time point (**fig. 3b**, RM ANOVA, F_timeXsex_ (5,125)=0.98, *p*=0.43, F_time_ (5,125)=30.58, *p*<0.0001, F_sex_ (1,25)=7.72, *p*=0.62). To further examine the time course of AIH in each sex and determine whether the amount of alcohol consumed correlated with mechanical sensitivity within each sex, we analyzed males and females separately. In male mice, the mechanical threshold averaged 1.45g prior to drinking. We found a significant interaction between time and alcohol treatment (**fig. 3b**, RM ANOVA males F_timeXtreatment_ (5,125)=4.70, *p*=0.0006, F_time_ (5,215)=8.59, *p*<0.0001; F_treatment_ (1,25)=37.90, *p*<0.0001). Multiple comparison testing showed that at week 4, alcohol drinking males had reduced mechanical thresholds averaging 1.17g that persisted to week 6 compared with the water drinking males, an overall decrease of 19.3% (Sidak’s multiple comparisons test between male alcohol and water drinkers, \**p*=0.01 Week 4, \*\*\*\**p*<0.0001 Week 5, \**p*=0.02 Week 6). There was no correlation between g/kg alcohol consumption and tactile threshold at Week 2 (development of AIH and the week with greatest alcohol consumption), Week 4 (earliest time point after AIH established) or Week 5 (establishment of AIH) (**fig. 3c-e**, simple linear regression Week 2 F(1,30)=1.07, *p*=0.31; Week 4 F(1,30)=0.004, *p*=0.95; Week 5 F(1,30)=0.07, *p*=0.80).

In female mice, mechanical thresholds averaged 1.44g prior to drinking. Similarly, we found a significant interaction between time and alcohol treatment (**fig. 3b**, RM ANOVA females F_timeXtreatment_ (5,120)=8.23, *p*<0.0001, F_time_ (5,120)=15.33, *p*<0.0001; F_treatment_ (1,24)=27.61, *p*<0.0001). Multiple comparisons testing showed that at week 4, alcohol drinking females had reduced mechanical thresholds averaging 1.03g that persisted to week 6 compared with water drinking females, an overall decrease of 28.5% (Sidak’s multiple comparisons test between female alcohol and water drinkers, \*\*\*\**p*<0.0001 for Weeks 4, 5 and 6). There was also no correlation between the g/kg alcohol consumption and tactile threshold for female mice at Weeks 2, 4 or 5 (**fig. 3c-e**, simple linear regression Week 2 F(1,27)=0.46, p=0.51; Week 4 F(1,27)=1.56. *p*=0.22; Week 5 F(1,27)=0.20, *p*=0.66).

There was no correlation between body weight and paw withdrawal threshold at Week 5 for either sex (**fig. 3f**, simple linear regression males F(1,30)=0.143, *p*=0.71; females F(1,27)=0.568, *p*=0.46). These data indicate that neither the time course nor magnitude of AIH are correlated to the amount of alcohol consumed or the weights of the animals.

Cold sensitivity was evaluated before alcohol exposure and after the development of AIH at Week 8 of CIA-2BC. Interestingly, there was a clear sex effect as we found that females, but not males, exhibited alcohol-induced cold hypersensitivity (**fig. 4**). There was a time by sex by alcohol drinking group interaction (three-way RM ANOVA F_timeXsexXdrinking condition_ (1,27)=4.34, *p*=0.04, F_sexXdrinking condition_ (1,27)=1.71, *p*=0.20, F_timeXdrinking condition_ (1,27)=4.43, *p*=0.04, F_timeXsex_ (1,27)=5.05, *p*=0.03, F_drinking condition_ (1,27)=4.97, *p*=0.03, F_sex_ (1,27)=29.89, *p*<0.0001, F_time_ (1,27)=50.71, *p*<0.0001). Multiple comparisons testing revealed that in male mice, no significant differences in cold sensitivity were observed between alcohol-and water-drinking groups at either time point (**fig. 4**, Sidak’s multiple comparisons test, week 0 *p*>0.99, week 8 *p*>0.99, between male alcohol and water drinking mice). However, alcohol-drinking female mice developed significantly greater cold sensitivity after week 8 compared to female water drinkers (**fig. 4**, Sidak’s multiple comparisons test at week 8, \*\**p*=0.005 female alcohol vs water) and compared to male alcohol drinkers (**fig. 4**, Sidak’s multiple comparisons test at week 8, \*\*\*\**p*<0.0001 alcohol female vs alcohol male). These data suggest that alcohol-induced cold hypersensitivity is sex-dependent, emerging in females, but not in males.

**Figure 4.**
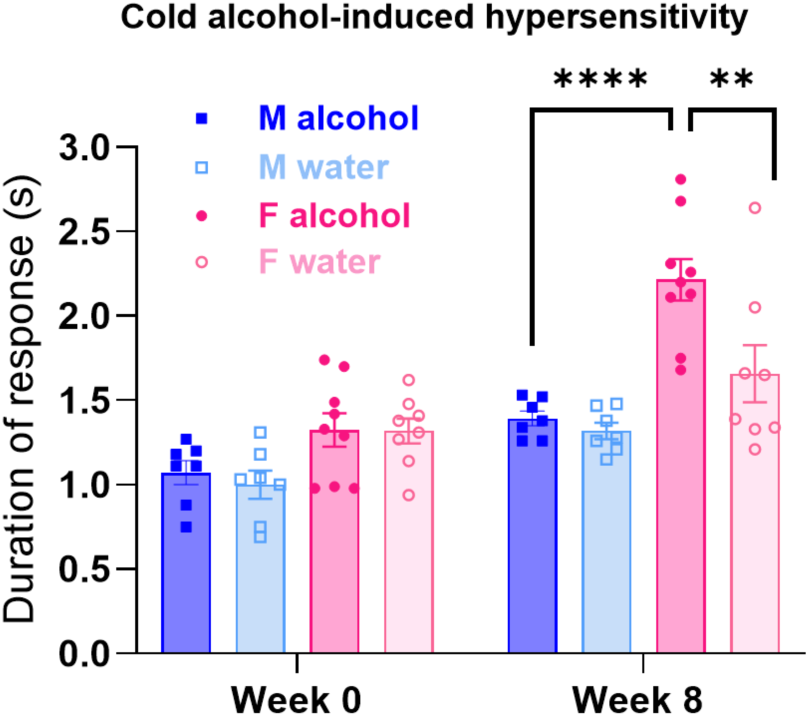
Chronic alcohol consumption produces alcohol-induced cold hypersensitivity in females only. Cold sensitivity was observed in female, but not male, mice after 8 weeks of drinking, Sidak’s multiple comparisons test between groups Week 8 \*\**p*<0.01, \*\*\*\**p*<0.0001. Males *n*=6-7/group, females *n*=8-9/group.

### 3.2 Chronic intermittent alcohol delays resolution of capsaicin-induced mechanical hypersensitivity

To investigate the role of voluntary alcohol consumption on recovery from acute inflammatory pain, mechanical thresholds were assessed following hind paw intradermal capsaicin injection in male and female mice that underwent chronic CIA-2BC. Capsaicin was administered during week 8 of CIA-2BC after mechanical AIH had emerged in both sexes (**fig. 5a**). Recovery was defined as the time point when the post-injection mechanical thresholds were significantly different from the +2 hour post-injection threshold (which was the time point of greatest sensitivity), indicating the mice were no longer hypersensitive. There were no significant differences in mechanical thresholds between the male and female alcohol-drinking groups following capsaicin, (RM ANOVA, no significant time by sex interactions, no main effects of time or main effects of sex, all *p*>0.05), which indicates that voluntary alcohol consumption does not affect the magnitude of hypersensitivity produced by capsaicin.

**Figure 5.**
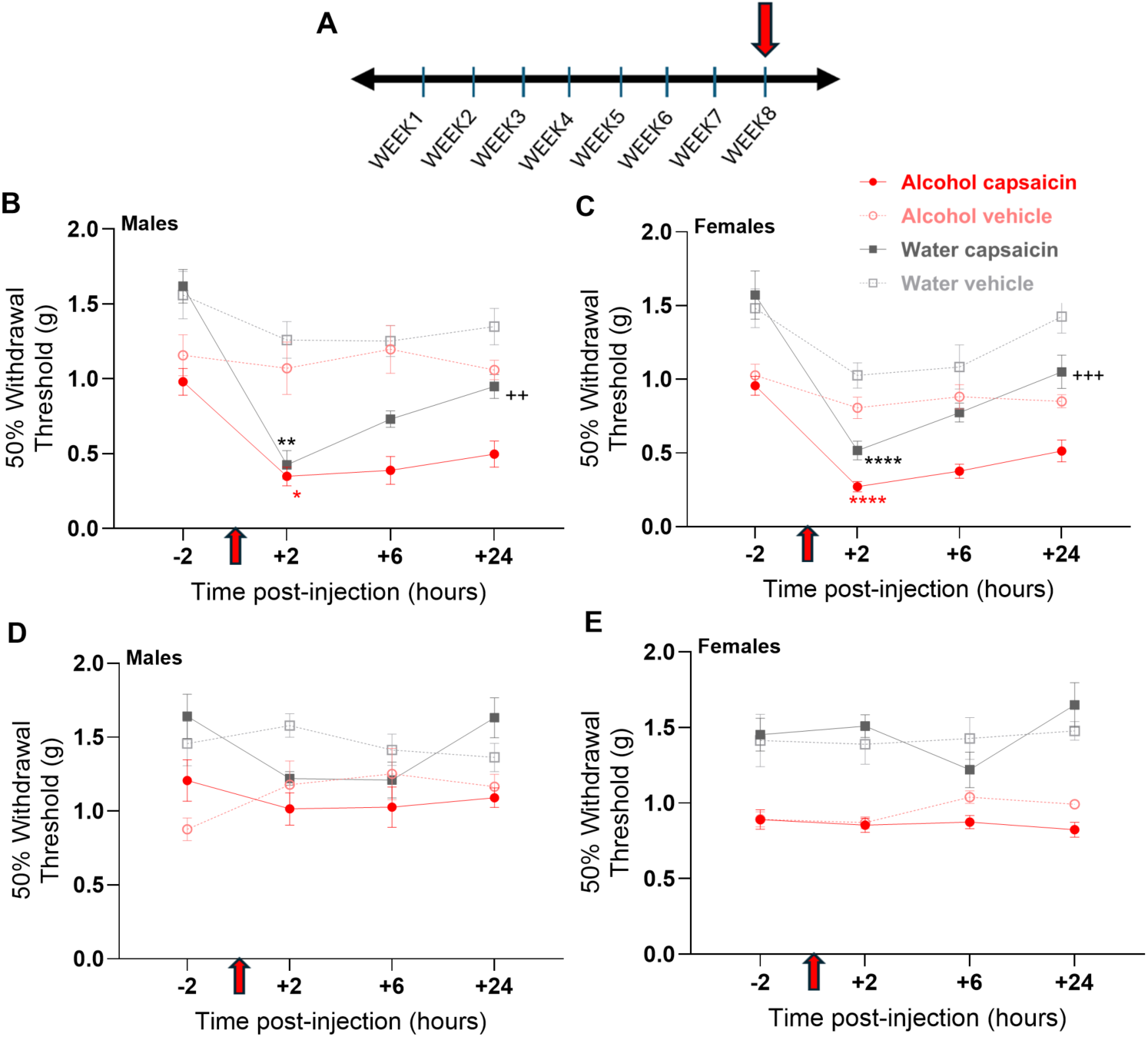
Chronic alcohol consumption delays resolution of mechanical hypersensitivity induced by capsaicin. Mechanical thresholds following hind paw intradermal capsaicin (30μg) injection in male (*n*=7-8/group) and female mice (*n*=6-7/group). **(A)** Red arrows indicate capsaicin or vehicle hind paw injection, which occurred after 8 weeks of CIA. Capsaicin significantly lowered tactile thresholds in the ipsilateral paw of **(B)** males and **(C)** females regardless of drinking group assignment (Sidak’s multiple comparisons test \**p*<0.05,\*\**p*<0.01,\*\*\*\**p*<0.0001 compared with -2hr within alcohol-or water-drinking groups). While water-drinking male and female mice had recovered to normal thresholds by 24 hours, hypersensitivity persisted in alcohol drinking mice (Sidak’s multiple comparisons test ++*p*<0.01 and +++*p*<0.001 compared with +2hr within alcohol-or water-drinking groups). Intradermal capsaicin had no significant effect on the non-injected paw in alcohol-or water-drinking group in **(D)** males or **(E)** females (*p*>0.05, Sidak’s multiple comparisons test).

Capsaicin-induced hypersensitivity normally resolves in 24-48 hours; we hypothesized chronic alcohol consumption would delay resolution of capsaicin-induced hypersensitivity. To further examine the time course and effect of alcohol consumption versus water consumption on capsaicin sensitivity, we examined males and females separately. In males, capsaicin reduced thresholds in the ipsilateral paw of both the alcohol-and water-drinking groups (**fig. 5b**, RM ANOVA, capsaicin F_drinking conditionXtime_ (3,39)=3.35, *p*=0.03, F_drinking condition_ (1,13)=58.04, *p*<0.0001, F_time_ (3,39)=38.11, *p*<0.0001; vehicle F_drinking conditionXtime_ (3,36)=0.70, *p*=0.56, F_drinking condition_ (1, 12)=4.38, *p*=0.06, F_time_ (3,36)=0.90, *p*=0.45). At 2 hours post-capsaicin injection, there were no differences in thresholds between alcohol-and water-drinking mice (Sidak’s multiple comparisons test, *p*=0.95 +2hr), suggesting that alcohol consumption has no effect on the magnitude of acute capsaicin-induced hypersensitivity. Mechanical thresholds significantly recovered after 24 hours in water-drinking capsaicin-injected mice (**fig. 5b**, Sidak’s multiple comparisons test, \*\**p*=0.008 +2hr vs. +24hr). However, alcohol-drinking capsaicin-injected mice failed to recover after 24 hours (**fig. 5b**, Sidak’s multiple comparisons test, *p*=0.75 +2hr vs. +24hr). This suggests chronic alcohol consumption prolongs hypersensitivity after acute inflammation.

In females, capsaicin reduced thresholds in the ipsilateral paw after 2 hours compared to baseline (**fig. 5c**, RM ANOVA, capsaicin F_drinking conditionXtime_ (3,33)=1.84, *p*=0.16, F_drinking condition_ (1,11)=68.26, *p*<0.0001, F_time_ (3,33)=38.80, *p*<0.0001; vehicle F_groupXtime_ (3,30)=1.72, *p*=0.18, F_drinking condition_ (1,10)=24.58, *p*=0.0006, F_time_ (3,30)=4.85, *p*=0.007). As in males, alcohol consumption had no effect on the magnitude of acute capsaicin-induced hypersensitivity between alcohol- and water-drinking mice (**fig. 5c**, Sidak’s multiple comparisons test, *p*=0.16 at +2hr). In water-drinking mice, this mechanical sensitivity was significantly reversed by 24 hours post-injection (**fig. 5c**, Sidak’s multiple comparisons test, \*\*\**p*=0.0009 +2hr vs. +24hr). However, recovery was not observed in alcohol-drinking mice (**fig. 5c**, Sidak’s multiple comparisons, *p*=0.24 +2hr vs. +24hr), who retained a similar degree of mechanical hypersensitivity as at 2 hours post-injection.

As expected, capsaicin had no effect on mechanical thresholds of the contralateral paw in either sex; however, there was a trend for a significant difference in mechanical thresholds in the contralateral paw in the control males (Sidak’s multiple comparisons test at -2hr vs. +2hr, *p*=0.76 alcohol-males, *p*=0.08 control-males, *p*=0.99 alcohol-females, *p*=0.99 control-females). There was a main effect of alcohol drinking in the capsaicin and vehicle treated groups, in both males and females, which indicated that AIH persisted in the non-injected hind paw (**fig. 5d**, males RM ANOVA, capsaicin F_drinking conditionXtime_ (3,39)=1.23, *p*=0.31, F_time_(3,39)=4.15, *p*=0.01, F_drinking condition_ (1,13)=12.34, *p*=0.004; vehicle F_drinking conditionXtime_ (3,36)=1.23, *p*=0.31, F_time_ (3,36)=1.16, *p*=0.36, F_drinking condition_ (1,12)=17.17, *p*=0.001; **fig. 5e**, females RM ANOVA, capsaicin F_drinking conditionXtime_ (3,33)=2.34, *p*=0.08, F_drinking condition_ (1,11)=178.4, *p*<0.0001, F_time_ (3,33)=1.56, *p*=0.22; vehicle F_timeXdrinking condition_ (3,30)=0.27, *p*=0.85, F_drinking condition_ (1,10)=27.22, *p*=0.0004, F_time_ (3,30)=0.82, *p*=0.50).

### 3.3 Chronic intermittent alcohol delays resolution of mechanical hypersensitivity induced by nerve injury

To investigate how CIA-2BC consumption affects persistent pain, we performed a sciatic nerve crush or sham surgery on mice 6 weeks after the initiation of drinking (**fig. 6a**). We chose this model of nerve injury because it induces transient nerve injury-related mechanical and thermal hypersensitivity [42, 48]. Nerve injury-induced mechanical hypersensitivity resolves after 3-5 weeks in healthy adult mice while heat hypersensitivity resolves in about 4 weeks in healthy adult rats [48]. We hypothesized chronic alcohol consumption would delay resolution of nerve injury-induced hypersensitivity in both sexes. For this analysis, recovery was defined as the time point when the post-injury tactile threshold was significantly different from the week 1 post-injury threshold. Week 2 data was excluded since mice were not tested this week due to technical issues.

**Figure 6.**
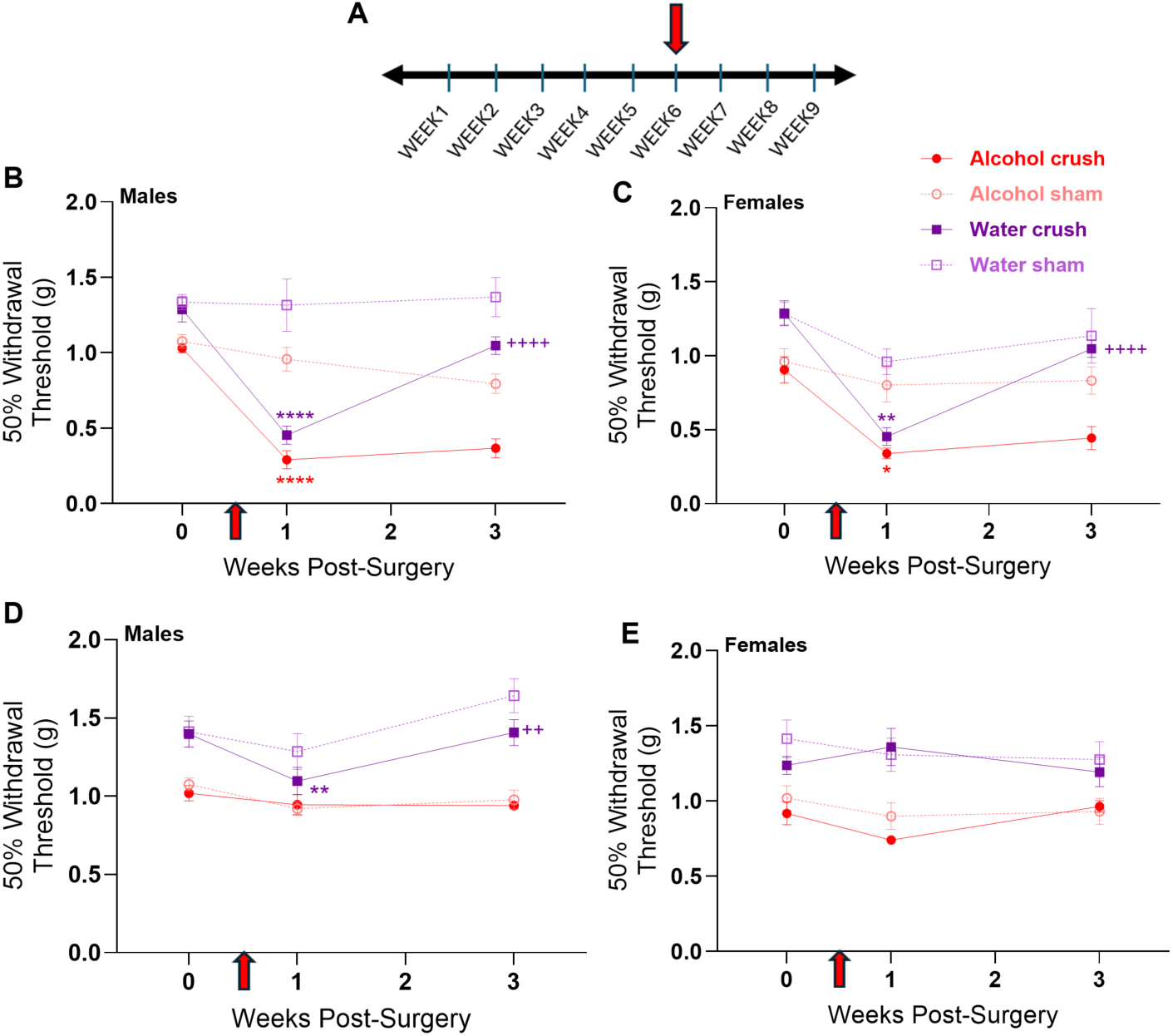
Chronic intermittent alcohol delays resolution of mechanical hypersensitivity induced by transient nerve injury. **(A)** Red arrows indicate sciatic crush/sham surgery was performed after 6 weeks of CIA, with CIA continuing for a total of 9 weeks. Mechanical thresholds in **(B, C)** ipsilateral injured and **(D, E)** contralateral uninjured hind paws across all weeks of the experiment (\**p*<0.05,\*\**p*<0.01,\*\*\*\**p*<0.0001 compared within group to pre-surgery week 0, ++*p*<0.01 and ++++*p*<0.0001 compared within group to post-surgery week 1, Sidak’s multiple comparisons test). Male and female data had no significant differences between thresholds of the same treatment group.

Similar to the results observed with capsaicin injection, there were no significant differences in mechanical thresholds between the male and female alcohol-drinking groups that had the nerve crush injury at all three time points measured, indicating that injury-induced mechanical thresholds were not affected by the differences in voluntary alcohol consumption (RM ANOVA alcohol-crush F_timeXsex_ (2,28)=3.03, *p*=0.06, F_time_ (2,28)=125.9, *p*<0.0001, F_sex_ (1,14)=6.5*10^-5^, *p*=0.99). To further examine the time course and effect of alcohol consumption versus water consumption on nerve injury-induced sensitivity, we examined males and females separately.

In male mice, as expected, crush injury produced mechanical hypersensitivity in the ipsilateral paw the week following injury regardless of drinking group (**fig. 6b**, RM ANOVA, crush F_timeXdrinking condition_ (2,30)=14.02, *p*<0.0001, F_time_ (2,30)=136.6, *p*<0.0001, F_drinking condition_ (1,15)=40.84, *p*<0.0001). The week following injury, water- and alcohol-drinking groups both showed reduced thresholds with no significant difference between the drinking conditions (**fig. 6b**, Sidak’s multiple comparisons test, *p*=0.28), indicating that chronic alcohol consumption does not affect the magnitude of nerve injury-induced hypersensitivity. While nerve injured mice drinking water recovered to their pre-surgical thresholds at 3 weeks post-injury, nerve injured mice drinking alcohol failed to recover at this time point (**fig. 6b**, Sidak’s multiple comparisons test, water-crush \*\*\*\**p*=0.0001 week 1 vs. week 3, alcohol-crush *p*=0.65 week 1 vs. week 3). This suggests chronic alcohol consumption prolongs mechanical hypersensitivity after nerve injury. In male mice that underwent sham surgery, we found a main effect of alcohol group only, indicating that sham surgery had no impact on thresholds, and that alcohol drinking lowered thresholds in the ipsilateral paw indicative of AIH (**fig. 6b**, RM ANOVA, sham F_timeXdrinking condition_ (2,28)=1.80, *p*=0.18, F_time_ (2,28)=1.07, *p*=0.36, F_drinking condition_ (1,14)=14.36, *p*=0.03).

In females, we also observed that crush injury produced mechanical hypersensitivity in the ipsilateral paw the week following injury regardless of drinking condition (**fig. 6c**, RM ANOVA, crush F_timeXdrinking condition_ (2,24)=10.61, *p*=0.001, F_time_(2,24)=88.51, *p*<0.0001, F_drinking condition_ (1,12)=22.55, *p*=0.001). Multiple comparisons tests showed that one week following injury, the mechanical thresholds were not significantly different between the water-drinking versus the alcohol-drinking females (**fig. 6c**, Sidak’s multiple comparisons test, *p*=0.25), again indicating that chronic alcohol consumption does not affect the magnitude of nerve injury-induced hypersensitivity. Water-drinking female mice recovered from nerve injury-induced hypersensitivity after 3 weeks, while alcohol-drinking mice showed no improvement (**fig. 6c**, Sidak’s multiple comparisons test, water-crush \*\*\*\**p*=0.0001 week 1 vs. week 3, alcohol-crush *p*=0.45 week 1 vs. week 3). These data show that in females, chronic alcohol consumption also prolongs mechanical hypersensitivity after nerve injury. In female mice that underwent sham surgery, we also found a main effect of alcohol group only, indicating that sham surgery had no impact on thresholds, and that alcohol drinking lowered thresholds in the ipsilateral paw indicative of AIH (**fig. 6c**, RM ANOVA, sham F_timeXdrinking condition_ (2,20)=0.32, *p*=0.73, F_time_(2,20)=2.31, *p*=0.12, F_drinking condition_ (1,10)=7.82, *p*=0.02).

For the contralateral paw, we found either an interaction between time and drinking condition or a main effect of drinking group in male and female mice (**fig. 6d**, RM ANOVA, crush F_timeXdrinking condition_ (2,30)=3.94, *p*=0.05, F_drinking condition_ (1,15)=29.65, *p*<0.0001, F_time_(2,30)=5.00, *p*=0.01; sham F_timeXdrinking condition_ (2,30)=2.55, *p*=0.10, F_drinking condition_ (1,15)=48.80, *p*<0.0001, F_time_(2,30)=3.46, *p*=0.04) and females (**fig. 6e**, RM ANOVA, crush F_timeXdrinking condition_ (2,24)=4.04, *p*=0.03, F_drinking condition_ (1,12)=29.92, *p*=0.0001, F_time_(2,24)=0.11, *p*=0.90; sham F_timeXdrinking condition_ (2,20)=0.05, *p*=0.95, F_drinking condition_ (1,10)=20.77, *p*=0.001, F_time_(2,20)=0.79, *p*=0.47). Together with the data on the ipsilateral paw, these findings indicated that mechanical AIH affects both hind paws and that it persisted through the entire duration of the drinking paradigm. In the contralateral paw of the nerve injured mice, we found no significant differences in thresholds pre-vs. post-injury (week 0 versus week 1) in the alcohol-crush males, alcohol-crush females or the water-crush females (**fig. 6d,e**, Sidak’s multiple comparisons test week 0 vs. week 1, alcohol-crush males *p*=0.79, alcohol-crush females *p*=0.25, and water-crush females *p*=0.57). However, we found an unexpected difference in contralateral paw thresholds in the water-crush males (**fig. 6d**, week 0 vs. week 1, \*\**p*=0.008).

### 3.4 Female mice drink more than male mice regardless of injury

We then examined alcohol consumption to investigate the effect of nerve injury on alcohol intake. There was no interaction between time, sex, and surgery treatment for average alcohol consumption (**fig. 7a**, F_timeXsexXsurgery treatment_ (8,208)=1.60, *p*=0.13, F_timeXsex_ (8,208)=2.18, *p*=0.03, F_timeXsurgery treatment_ (8,208)=0.88, *p*=0.53, F_sexXsurgery treatment_ (1,26)=0.98, *p*=0.33, F_time_ (8,208)=2.74, *p*=0.007, F_sex_ (1,26) = 65.30, *p*<0.0001, F_surgery treatment_ (1,26)=2.61, *p*=0.12). Similarly, there was no significant interaction for alcohol preference (**fig. 7b**, F_timeXsexXsurgery treatment_ (8,208)=1.44, *p*=0.18, F_timeXsex_ (8,208)=3.31, *p*=0.001, F_timeXsurgery treatment_ (8,208)=0.71, *p*=0.69, F_sexXsurgery treatment_ (1,26)=0.04, *p*=0.85, F_time_ (8,208)=3.58, *p*=0.0007, F_sex_ (1,26) = 3.30, *p*=0.08, F_surgery treatment_ (1,26)=1.43, *p*=0.24). There was no interaction or main effect of the surgery treatment (crush vs. sham) on alcohol consumption or preference, indicating that the crush injury had no significant impact on alcohol consumption or preference. The data was then consolidated such that crush and sham groups were combined for each sex. Analysis of the consolidated data revealed an interaction between time and sex for average alcohol consumption (**fig. 7c**, F_timeXsex_ (8,224)=2.18, *p*=0.03, F_time_ (8,224)=2.73, *p*=0.007, F_sex_ (1,28)=61.63, *p*<0.0001) and alcohol preference (**fig. 7d**, F_timeXsex_ (8,224)=3.12, *p*=0.002, F_time_ (8,224)=3.55, *p*=0.0007, F_sex_ (1,28)=3.38, *p*=0.08), with Sidak’s multiple comparisons tests showing that females have higher alcohol consumption and preference compared with males at multiple time points in the CIA procedure. These findings are similar to the increased alcohol consumption and preference observed in females produced by CIA alone (**fig. 2**). These data also suggest that nerve injury-induced hypersensitivity has no effect on alcohol consumption or preference in the CIA-2BC.

**Figure 7.**
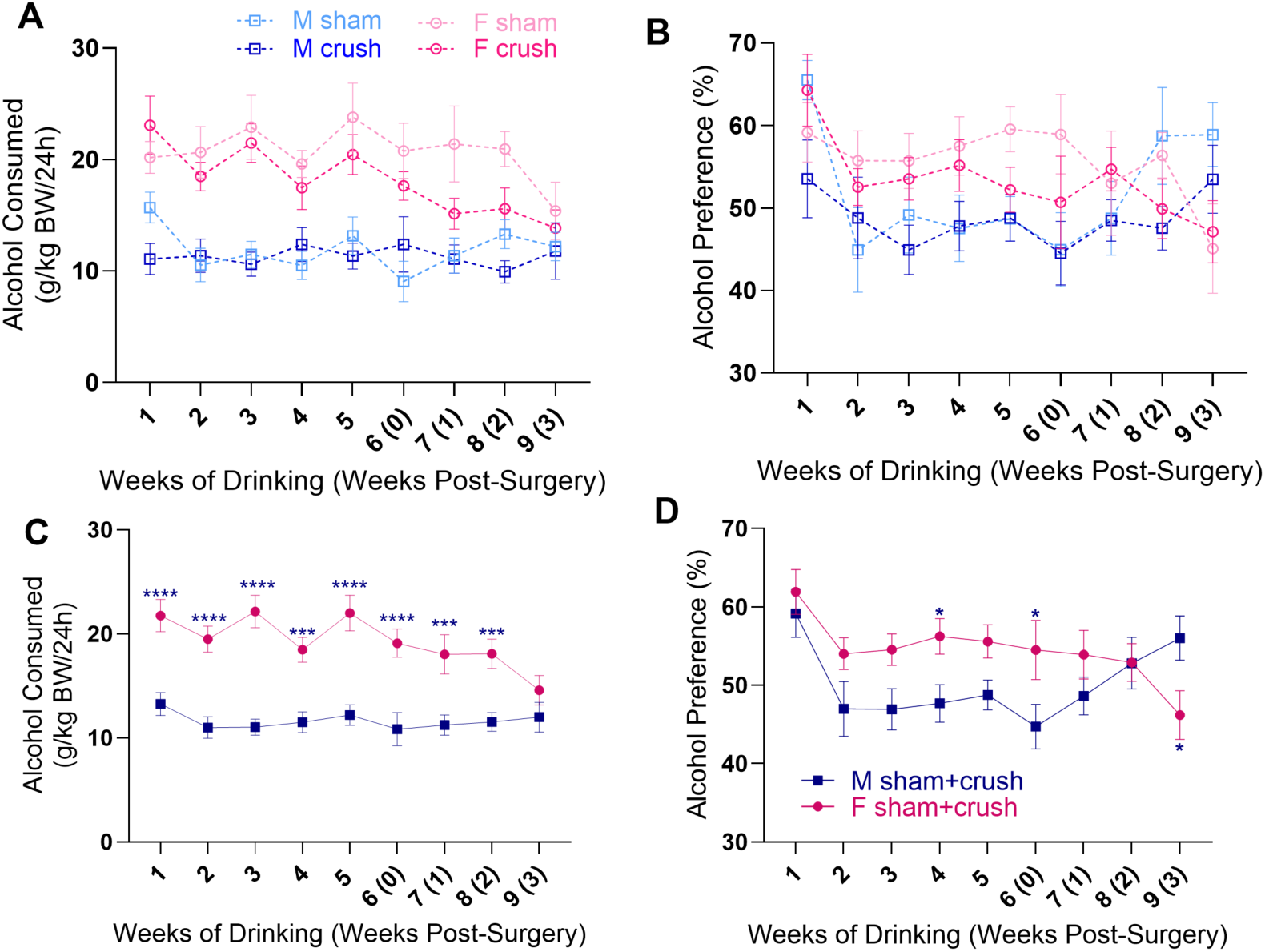
Female mice voluntarily drink more alcohol than male mice regardless of injury. **(A)** Average alcohol consumed per day measured relative to body weight and **(B)** % preference for the alcohol bottle for males (*n*=8-9/group) and females (*n*=6-7/group). There was no effect of injury on alcohol consumption (3-way RM ANOVA *p*>0.05). **(C)** Alcohol consumption and **(D)** preference shown with the sham and crush groups combined within sex. Female mice consumed more alcohol (g/kg/24h) and had a higher preference for the alcohol bottle compared with male mice (Sidak’s multiple comparisons test between male sham + crush vs. female sham + crush at each time point, \**p*<0.05, \*\*\**p*<0.001, \*\*\*\**p*<0.0001).

## 4. Discussion

Our data indicate that voluntary chronic alcohol consumption increases mechanical sensitivity and extends recovery time following acute and persistent injury. Neither the difference in the amount of alcohol consumed between males and females, nor the amount of alcohol consumed within sex, influence the magnitude of alcohol-induced hypersensitivity. These data suggest the effects of alcohol on mechanical sensitivity are not dose-dependent but rather may be associated with a threshold level of alcohol consumption or duration of alcohol consumption. Moreover, differences in the amount of alcohol consumed between males and females did not affect the magnitude of mechanical hypersensitivity produced by acute or persistent injury, nor the time course of alcohol-induced prolongation of recovery from injury. CIA also increases cold sensitivity in females and not in males, illustrating sex differences in the mechanisms of alcohol-induced sensitivity.

To our knowledge, the present study is the first to investigate how chronic voluntary alcohol consumption affects recovery from mechanical hypersensitivity in different pain models by monitoring mechanical thresholds over time. We found that chronic alcohol drinking delays recovery from capsaicin-induced mechanical hypersensitivity and transient nerve injury-induced mechanical hypersensitivity. These results suggest that chronic voluntary alcohol consumption may facilitate the transition to chronic pain by prolonging hypersensitivity and delaying recovery from injury. The mechanisms underlying this delayed recovery could be either peripheral, central, or a combination of both. Repeated exposure of free nerve endings to alcohol could be neurotoxic (in the case of small fiber neuropathies) or increase inflammation and oxidative stress, “priming” nociceptors to be more excitable. Alcohol could also sensitize dorsal root ganglia (DRG) neurons by promoting inflammation or alcohol-induced changes in gene expression that produce heightened excitability. Centrally, alcohol in the spinal cord may enhance NMDAR signaling while reducing GABAergic signaling in the dorsal horn, altering normal inhibitory tone modulated in the dorsal horn. Lastly, chronic alcohol exposure may disrupt brain circuits, especially limbic regions and descending pain modulatory circuits that contribute to pain facilitation and inhibition.

Consistent with other rodent studies [34–36], we found female mice consume more alcohol than male mice. In both sexes, nerve injury-induced hypersensitivity did not affect alcohol consumption in this intermittent access protocol measured via daily bottle weights. It is possible that gross daily bottle weight may be unable to capture potential changes in alcohol drinking behavior in mice. Additionally, a ceiling effect on alcohol intake in our model may have masked injury-induced increases in alcohol consumption, as mice were already drinking at high levels. For example, other studies using the CIA-2BC paradigm report average 24h intakes of 15-20 g/kg in males and 25-30 g/kg in females [35, 37], suggesting a potential ceiling effect around 20 and 30 g/kg for males and females, respectively. Our mice were drinking ∼15 g/kg for males and ∼25 g/kg for females in the nerve crush experiment, indicating intake was already near maximal levels.

The lack of effect of nerve injury-induced hypersensitivity on alcohol intake contrasts with clinical literature, which suggests individuals with pain drink more alcohol to self-medicate pain symptoms [6–9], and with preclinical evidence of anti-hyperalgesic and anti-allodynic effects of acute alcohol [49, 50]. However, the preclinical evidence for the relationship between chronic alcohol consumption and persistent pain is contradictory. In male and female rats with prior alcohol exposure in a CIA-2BC procedure, hind paw injections of complete Freund adjuvant did not increase or decrease intake for 20% alcohol [51].

In alcohol-naïve rodents, induction of persistent pain had highly variable effects on chronic alcohol consumption. Persistent inflammatory pain was associated with increased alcohol intake in male mice only [35] but suppressed intake in male and female rats [52] in a continuous access 2BC paradigm. In a surgical incision model of postoperative pain, male but not female mice consumed more alcohol (continuous access 2BC) [53], whereas binge drinking was decreased in male but not female mice with spared nerve injury [54]. These variable observations may stem from differences in alcohol exposure paradigms, pain models, and the presence or absence of prior alcohol exposure; taken together, they reflect the complexity of the chronic alcohol-pain relationship. It is possible that prolonged alcohol exposure disrupts the neural substrates that mediate the acute analgesic effects of alcohol.

Our data replicated findings that CIA consumption produces alcohol-induced mechanical hypersensitivity, in agreement with prior clinical and preclinical reports. Moreover, our data showed that CIA consumption produces alcohol-induced cold hypersensitivity in female but not male mice. We also found no direct correlation between paw withdrawal thresholds and amount of alcohol consumed at weeks 2-5 of CIA, which correspond to the period of establishment of alcohol preference and AIH. The emergence of AIH in our model aligns with previous human and rodent studies demonstrating that chronic intermittent alcohol exposure – via vapor inhalation or 2BC – leads to mechanical hypersensitivity [19, 21–25]. These effects are often observed during protracted withdrawal or following repeated acute withdrawal episodes within intermittent exposure paradigms. In humans, alcohol-related neuropathy is a common consequence of chronic alcohol consumption [55]. Small fiber peripheral neuropathy is among the most frequent neurologic symptoms of AUD, often producing greater loss of sensation in the legs than in the arms [56]. Although the precise mechanisms underlying AIH remain unclear, AIH could result from prolonged neuroinflammation leading to peripheral and/or central sensitization. One study observed that after CIA in alcohol-dependent mice, there was increased microglial activation and elevated levels of pro-inflammatory IL-6 in the spinal cord [25]. Alcohol potentiates nociceptor responses *in vitro* [57], but more research is needed regarding the effects of chronic alcohol consumption on primary afferents *in vivo*. Disinhibition of the descending pain inhibition pathway could also contribute to AIH. Both potential mechanisms would amplify responses in ascending pain pathways and result in greater sensitivity.

The effects of alcohol on cold sensitivity remain largely understudied in rodent and human subjects as most research emphasizes heat sensitivity [19]. Our study is the first to assess cold sensitivity during chronic alcohol consumption. Cold sensitivity was significantly increased in female alcohol drinkers compared to female water drinkers and male alcohol drinkers, indicating a sex-specific effect of alcohol on cold sensitivity. These findings are consistent with broader patterns of female-specific pain sensitivity observed preclinically and clinically. Rodent studies have shown greater mechanical hypersensitivity in females in chronic pain models [58–60] and sex-specific emergence of heat hypersensitivity after chronic alcohol exposure [19]. Clinically, many chronic pain conditions – fibromyalgia, migraine, painful temporomandibular disorders, and chronic pelvic pain – are more prevalent in women. It is therefore possible that sex differences in alcohol-induced cold sensitivity reflect differential modulation of peripheral or central pain-processing mechanisms that contribute to a higher prevalence of chronic pain in females.

There are some limitations to consider with this study. First, social isolation may produce stress and exacerbate alcohol-induced hypersensitivity, considering some studies suggest social isolation in mice contributes to lower tactile thresholds following nerve injury thresholds [61, 62]. While mice were singly housed to accurately record daily alcohol consumption, group housing could provide a more natural social environment. Second, assessing noxious heat and cold sensitivities would have provided more insight into the multimodality of alcohol-induced hypersensitivity following inflammation and nerve injury. Heat sensitivity is heightened following chronic alcohol exposure [19]; thus, it will need to be assessed in our model in the future. Finally, we did not extend the testing period long enough to fully monitor the recovery time course in the CIA mice exposed to capsaicin or sciatic nerve crush. This will be considered in future experiments.

In conclusion, we demonstrated that CIA induces mechanical and sex-specific cold hypersensitivity and prolongs mechanical hypersensitivity after acute inflammation and nerve injury. These findings highlight the detrimental effects of chronic alcohol consumption on recovery from tissue damage and pain modulation. From a clinical perspective, our results emphasize the need for targeted pain management strategies in patients with chronic alcohol use, as their altered pain processing may complicate recovery and increase the risk of persistent pain. Therefore, it is essential to investigate the spinal and supraspinal neural mechanisms involved in the complex, comorbid relationship between chronic alcohol use and persistent pain.

## CRediT authorship statement

**Lucy Vulchanova:** conceptualization, funding acquisition, methodology, writing – review and editing. **Anna M. Lee:** conceptualization, methodology, writing – review and editing. **Laura Stone:** conceptualization, methodology, writing – reviewing and editing. **Maureen Riedl:** investigation. **Rachel Schorn:** investigation, data curation, writing – original manuscript, reviewing and editing. All authors reviewed the manuscript.

